# The tell-tale hair: Effects of dam contact and hutch type on hair cortisol concentrations in female Holstein calves

**DOI:** 10.64898/2026.11.25.769259

**Authors:** Ina Klawes, Mischa Alexej Weiß, Marvin Suhr, Eva Wasserloos, Anita Seidel, Stéphanie Céline Hornburg

## Abstract

Early rearing conditions and the social environment have decisive influences on calves’ stress responses. The effects of prolonged postpartum cow–calf contact and preweaning housing conditions on hair cortisol concentration (HCC) in Holstein calves are not yet fully understood. This study aimed to understand how different postpartum contact times between calves and their dams and subsequent hutch types affect cortisol concentrations in calf hair. Calves were separated from their dams either within 12 h after birth (S) or after 24–48 h (L) and subsequently housed individually (S1, L1) or in pairs (S2, L2). Total cortisol concentration was measured in the growing hair of 35 female Holstein calves after weaning (88 ± 9 days) using an enzyme-linked immunosorbent assay (ELISA) on a commercial farm located at 53°55’54.8”N 9°55’47.8”E and 15 metres above mean sea level. The HCC of S2-calves was lower compared with S1-calves (*p* = 0.047). In contrast, no differences (*p* > 0.05) between hutch types were observed in the prolonged cow–calf contact. The results suggest that pair housing reduces HCC during the preweaning phase in calves with a short contact time with the dam after birth.

## 1. Introduction

Germany is the largest milk-producing country in the European Union, with approximately 32.2 million tons of raw cows’ milk delivered to dairies in 2024 [1]. Holstein cattle are the predominant dairy breed in Germany and are widely represented in European dairy production, making Holstein calves a biologically and economically relevant model for investigating the effects of commercial rearing practices. In many dairy systems, calves are separated from their dams shortly after birth, housed individually during the first weeks of life, and subsequently transferred to group housing before or around weaning [2, 3]. Early separation and individual housing have traditionally been justified by management, health, and economic considerations, including supervision of colostrum intake, reduced pathogen transmission, prevention of uncontrolled suckling, and availability of saleable milk [4–8]. Nevertheless, these practices have received increasing public and scientific attention because of their potential effects on calf welfare and social development [9–12].

Greater postpartum contact between a cow and calf and early social housing have therefore been investigated as alternative rearing strategies. Studies of prolonged cow–calf contact systems have not provided consistent evidence that immediate separation is required to protect calf or cow health [13]. Such systems may facilitate maternal bonding, suckling, play, and other species-specific behaviours, although their implementation can require substantial changes in farm infrastructure and management [5, 8, 14].

Pair housing may represent a more readily applicable strategy for providing calves with social contact during the preweaning period. Compared with individually housed calves, socially housed calves exhibit more developed social behaviour, greater behavioural flexibility, reduced fear of unfamiliar animals and situations, and improved adaptation to regrouping and feeding transitions [15–20]. However, most previous studies have focused on behavioural outcomes. The extent to which early maternal and social experiences influence longer-term physiological responses to subsequent management stressors remains insufficiently understood [20]. Cortisol concentrations in blood, saliva and urine are frequently used as indicators of short-term hypothalamic–pituitary– adrenal axis activity [21–26]. However, these measurements mainly reflect cortisol secretion over relatively short periods, and blood sampling or handling may itself affect the measured response [22].

Hair cortisol concentration provides a complementary measure because cortisol incorporated into growing hair can reflect cumulative exposure over periods of weeks to months [27–29]. Although hair cortisol is not suitable for detecting rapid fluctuations in cortisol secretion, it may be useful for evaluating integrated physiological responses to prolonged or repeated management challenges. This is particularly relevant during calf rearing, when separation, dietary transition, weaning, and regrouping may occur within a relatively short developmental period [30, 31].

The present study aimed to understand how the duration of postpartum cow–calf contact and subsequent preweaning housing affect the hair cortisol concentration of female Holstein calves. Calves were separated from their dams either within 12 h after birth (S) or after 24–48 h (L) and subsequently housed individually (S1, L1) or in pairs (S2, L2). We hypothesised that S2- and L2-calves would exhibit lower hair cortisol concentrations during later management transitions than S1- and L1- calves because early social contact may improve their capacity to adapt to weaning and regrouping. We also examined whether postpartum contact duration modified the physiological response to housing treatment. Total serum protein concentration and body weight gain were assessed as supporting indicators of colostrum transfer, health, and development during the experimental period.

## 2. Materials and Methods

### 2.1. Animals, management, and study design

The study was conducted at Kiel University’s experimental dairy farm, Karkendamm, in Bimöhlen, Germany (53°55’54.8”N 9°55’47.8”E, 15 m above mean sea level), located in the federal state of Schleswig-Holstein in northern Germany. In Schleswig-Holstein, the long-term mean annual temperature is 9.3 °C, with an average temperature of 10.0 °C in 2025. The long-term mean annual precipitation in Schleswig-Holstein is 812.8 mm, with 728.4 mm in 2025. The long-term mean annual sunshine duration is 1,657.4 h, with 1,847.9 h in 2025. Climate data were obtained from the German Meteorological Service (Deutscher Wetterdienst, DWD).

All experiments involving animal handling procedures were carried out with the permission of the Kiel University animal welfare officer and the Schleswig-Holstein local authority (MLLEV-SH 64-12/23), in accordance with the relevant institutional and national guidelines for the care and use of laboratory animals (German animal welfare law [TierSchG] and TierSchVersV, the national implementation of Directive 2010/63/EU). All animal handling and sample collection procedures were conducted by specifically trained personnel, with particular care taken to minimise stress and ensure animal welfare.

Female Holstein-Friesian calves (n = 35), born on the farm between August 2024 and May 2025, were included in this study. Only female calves were used to minimise sexual variation. At birth, the calves were allocated to four experimental groups based on postpartum cow–calf contact time and subsequent housing type. The groups were S1 and S2 for short contact (< 12 h) followed by single or pair housing, respectively, and L1 and L2 for prolonged contact (24 - 48 h), followed by single or pair housing, respectively (see Table 1). The experimental unit for the study was the calf. Because calves assigned to pair housing had to be matched with a partner born within 24 h, suitable pairs during the experimental period were limited; consequently, approximately twice as many calves were housed individually as those in pairs. Each calf’s experimental period began at its birth and lasted until approximately one week after weaning (88 ± 9 days of age).

**Table 1.** Definition of the experimental groups.

| **Abbreviation** | **Cow–Calf Contact Time *pp*** | **Housing Type** | **n** |
|---|---|---|---|
| S1 | Short contact (< 12 h) | Single hutch | 12 |
| L1 | Prolonged contact (24 – 48 h) | Single hutch | 13 |
| S2 | Short contact (< 12 h) | Pair hutch | 5 |
| L2 | Prolonged contact (24 – 48 h) | Pair hutch | 5 |

#### 2.1.1 Calving pens and separation of cow and calves

Approximately three to four days before their calculated calving dates, the cows were removed from the herd and housed individually. The pens were 3.80 × 3.85 m, fitted with feeding racks and water troughs, and bedded with straw. If necessary, the cows received calving assistance from farm staff or a veterinarian. During the cow–calf contact period (up to 48 h, depending on the experimental group) in individual calving pens, calves had free access to their dams’ udders for suckling, and neither teat buckets nor nipple bottles were provided.

The cows and calves were separated at milking time. The calves in experimental groups S1 and S2 were separated from their dams after a postpartum contact time of 12 h, and the calves in experimental groups L1 and L2 were separated after a postpartum contact time between 24 and 48 h. The cows were herded to the fresh-milking group, where they remained as milking cows. The calves were herded into disinfected and freshly straw-bedded calf hutches either alone or in pairs with calves of the same age (±1 day).

#### 2.1.2 Housing conditions in single and pair hutches

Figure 1 shows the dimensions of the single and pair hutches, including the floor area and the associated enclosure area defined by a fence, as specified by the manufacturer. The housing and management of calves are regulated in Sections 5–11 of the German Tierschud-Nudtierhaltungsverordnung (TierSchNudtV). The calf hutches were located approximately 200 m from the calving barn and were all covered by a roofed shelter. The hutches were adjacent, and the calves could see each other, regardless of the experimental group. The calves stayed in the hutches until 28 ± 5 days of age.

**Figure 1.**
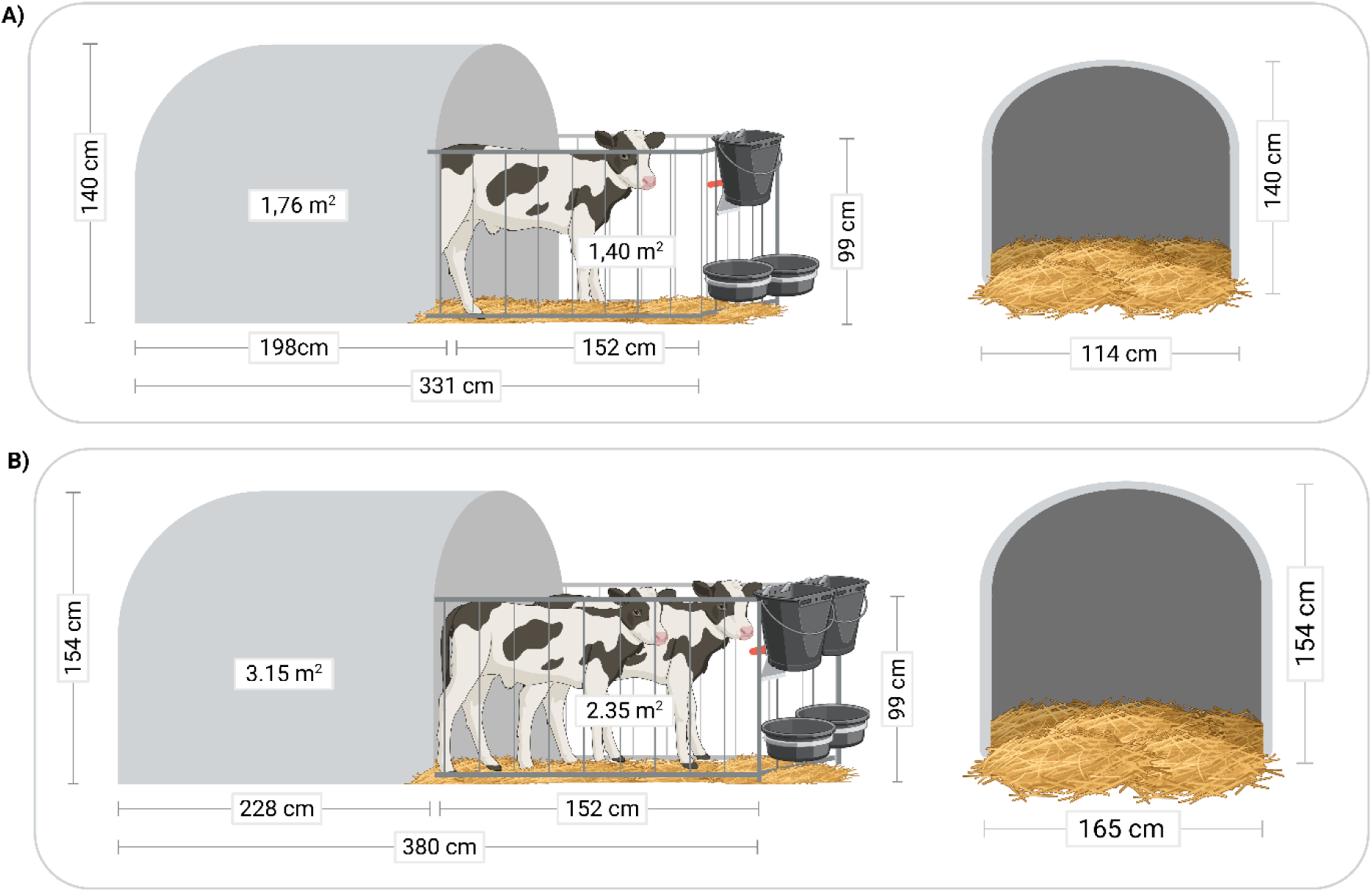
Schematic drawing of hutches. Both hutch types are bedded with straw and equipped with teat buckets at the front fence, as well as troughs for water and feed. The gates can be folded up for cleaning and disinfection after use, before new animals are housed in them. **A: Single hutch** (internal): 114 × 198 × 140 cm (width × depth × height); pen enclosure: 130 × 115 × 140 cm. The outer pen enclosure measures 114 × 152 × 99 cm (width × depth × height), corresponding to an enclosure area of 1.40 m². **B: Pair hutches** have the following dimensions: 165 × 228 × 154 cm (width × depth × height), providing approximately 3.15 m² of usable floor area. The outer pen enclosure measures 165 × 152 × 99 cm (width × depth × height), corresponding to an enclosure area of approximately 2.35 m². Created in BioRender. I. (2026). https://BioRender.com/i2uku27.

For the first three days of life, each calf was fed a minimum of 3 L colostrum twice daily via teat buckets placed in the hutch. The calves were carefully trained to drink from an individual teat bucket and, where possible, received colostrum from their own mothers. In instances where a mother cow failed to produce sufficient colostrum, the calf was provided with a mixture of colostrum from the cows that had given birth closest to its own birth. From four days of age onwards, the calves were fed up to eight litres of whole milk with whole milk enhancer (Farm-O-San StartFit, Trouw Nutrition, the Netherlands) from teat buckets twice daily. All calves received hay for ad libitum intake and concentrates (BAT Kälber Fit Plus FREE, BAT Agrar GmbH & Co. KG, Germany; Miarom Classic P, MIAVIT GmbH, Germany) with free access to water from water bowls.

#### 2.1.3 Housing conditions in group pens

At 28 ± 5 days of age, the calves were moved to straw-bedded group pens measuring 18 m², with up to five calves. Wherever possible, paired calves were kept together within the same group. The calves were housed in open pens with visual and tactile contact with other groups. Group pens were naturally lit and had artificial lighting from approximately 6 am to 5 pm during working hours. The calves were fed whole milk from plastic buckets with five teats (group feeders). Following relocation from the calf hutches to the group pens, the milk allowance was gradually reduced according to a predefined weekly weaning regimen (see Table 2) to transition the calves from milk to solid feed. The milk quantity was decreased stepwise from 8 l per day per calf over a ten-week period, leading to complete weaning by 12 weeks of age. For example, at 7 weeks of age, a group of five calves received 12.5 L whole milk per feeding via group feeder teat buckets.

**Table 2.** Milk feeding plan with amount of milk fed per calf during the weaning period.

| **Age (weeks)** | **Amount of Milk Fed (L)** | **Frequency per Day** |
|---|---|---|
| 5 | 4 | Twice |
| 6 | 3 | Twice |
| 7 | 2.5 | Twice |
| 8 | 2.5 | Twice |
| 9 | 2 | Twice |
| 10 | 1 | Twice |
| 11 | 1 | One time in the evening |
| 12 | 0 | Weaned |

Total mixed ration and concentrates (BAT Kälber Fit Plus FREE, BAT Agrar GmbH & Co. KG, Germany; Miarom Classic P, MIAVIT GmbH, Germany) were available ad libitum throughout the rearing period, and hay was provided ad libitum to encourage rumination and oral behaviour. All calves had free access to water bowls. After collecting the final sample, the calves were officially released from the animal experiment following a thorough examination by a veterinarian. Thereafter, they remained on the farm as conventional dairy cattle.

### 2.2. Sampling and data collection

On the 2nd day of age, whole blood samples were collected to monitor the colostrum supply, and the calves were shaved to allow for uniform hair growth during the experimental period. After weaning (88 ± 9 days of age), the calves were shaved again, and hair samples were collected for hair cortisol concentration analysis; see Figure 2 for a detailed overview of the study design and sampling time points. During herding into the calf hutches after cow–calf contact, the calves were weighed using a conventional calf scale (Holm & Laue Stationary Calf Scale WCB-000008, Holm & Laue, Germany). All animals were assessed three times weekly after feeding using an objective health check [32] to provide situation-specific recommendations for action when needed.

**Figure 2.**
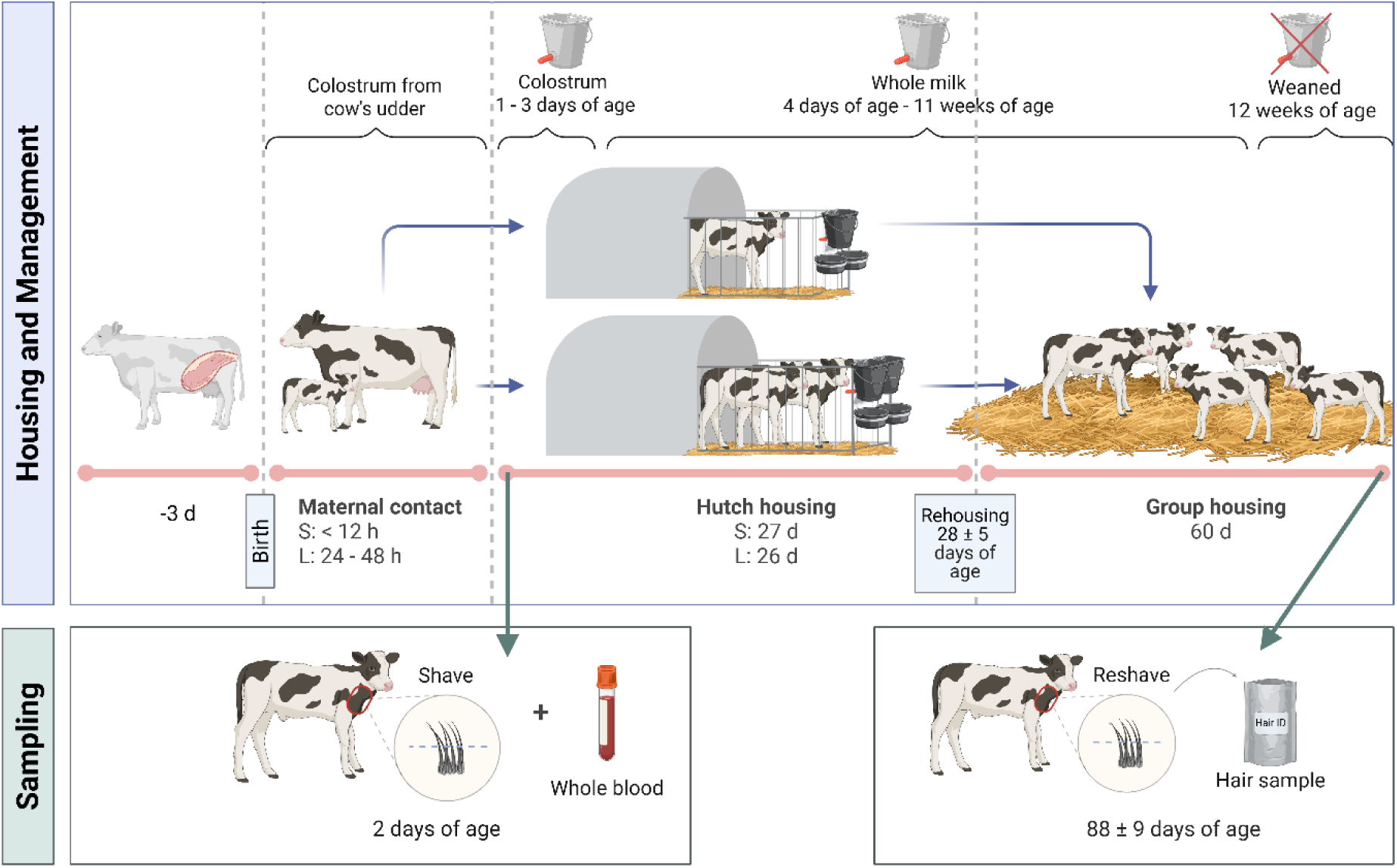
Overall workflow of the study: from pre-calving to postweaning. Three days before the calculated calving date, the cows were housed individually. After birth, female calves were separated from their dams after < 12 h or after 24 – 48 h and subsequently housed individually or in pair hutches. Whole blood samples were taken on the 2nd day of age, in line with the first shaving. For the first three days of age, the calves had colostrum from their dams. Subsequently, whole milk was administered until weaning. After 28 ± 5 days of age, the calves were housed in groups of up to five. One week after weaning (88 ± 9 days of age), hair samples were taken from the same spot where the first shaving occurred (reshaving). Created in BioRender. I. (2026). https://BioRender.com/i2uku27.

#### 2.2.1. Blood samples

As variations in cow–calf contact times precluded a standardised colostrum supply across experimental groups, the adequacy of passive transfer was evaluated by measuring serum Brix values via refractometry. Refractometry can be used to evaluate serum total protein and determine adequate passive transfer in calves [33–35]. Whole blood was collected from 2-day-old calves by jugular venipuncture using a 20-gauge, 1-inch hypodermic needle (BD Vacutainer Precision Glide, Becton Dickinson Co., Franklin Lakes, NJ) and placed into a sterile plastic Vacutainer tube without anticoagulant (BD Vacutainer, Becton Dickinson and Co.). The samples were transported under refrigeration to Kiel University.

#### 2.2.2. Hair samples

As recommended by Heimbürge et al. [36], hair samples were taken for cortisol determination using a minimally invasive sampling procedure by repeatedly (shave–reshave method) and carefully shearing the hair with an electric shearing machine (Series 3000 model BT3206/14; Philips, Amsterdam, The Netherlands). The calves were shaved on day 2 postpartum in the area where blood was sampled (Figure 3).

**Figure 3.**
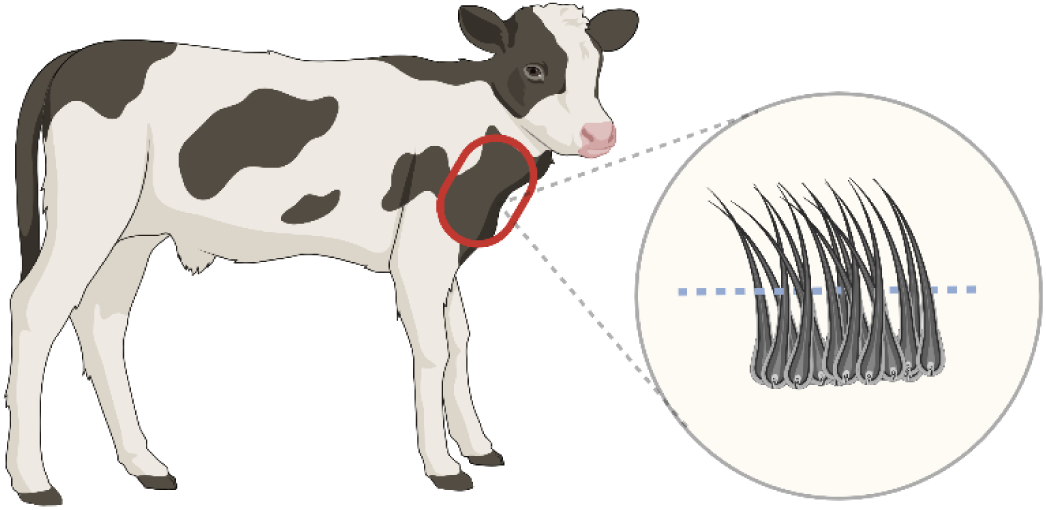
Sampling area for hair samples. On the 2nd day of age, a hair sample was taken from a calf’s neck with electric clippers.

Since the effect of hair colour on cortisol incorporation is controversially discussed, only black hair was sampled [29, 37, 38]. In the case of red-haired calves, red hair was sampled. Following the weaning phase, the same area was shorn again, and the hair that had grown back over the test period was used to evaluate long-term hair cortisol concentrations, as recommended [29]. The time delay from shave to reshave was considered to reflect cortisol incorporation according to the hair growth rate. The hair samples were labelled and brought to the laboratory in an envelope protected from light until further processing. The hair samples were dried; weighed; and, as a precaution, stored protected from light at room temperature until analysis [39, 40].

### 2.3. Laboratory procedures

#### 2.3.1. Serum total protein

Serum was separated from the whole blood samples by centrifugation at 3000×g for 5 min at room temperature (Heraeus Multifuge X3R, Thermo Fisher Scientific, Waltham, MA, USA) within 12 h after sampling. Serum Brix values (total protein) were then determined by duplicate measurement using a digital handheld refractometer (PAL-1, ATAGO Co., Ltd., Tokyo, Japan).

Handheld Brix refractometers are well-validated, standard side-test instruments for assessing serum protein and passive immunity status in calves. Multiple methodological studies have shown that handheld Brix readings correlate strongly with lab-determined total protein and serum IgG concentrations (r > 0.93; [35, 41, 42]). Accordingly, raw Brix values were utilized as a reliable surrogate to determine relative differences in serum protein concentrations across the experimental calf groups.

#### 2.3.2. Hair cortisol analysis by ELISA

The cortisol concentrations in the hair samples were analysed using a commercially available ELISA kit for quantifying free cortisol in saliva [27, 36, 43, 44]. Extraction and analysis of hair cortisol was performed using the procedures described by Meyer et al. [45] and Heimbürge et al. [29]. Each hair sample was placed in a 15 mL polypropylene tube with a screw cap, and 5 mL of isopropanol was added. The tube was gently mixed on a rotator at room temperature (RT) for 3 minutes, and after decanting, the washing step was repeated once. Then, the samples were dried for 2–3 days in clean, covered dishes at RT. To prepare the samples for grinding, 60 mg of dried hair was transferred from each sample into a 2 mL lysing tube containing six stainless steel grinding balls (Lysing Matrix S, MP Biomedicals). The samples were ground using a bead-based tissue homogeniser (FastPrep-24™ 5G, MP Biomedicals, USA) at a speed of 5.0 m/s for six cycles of 30 seconds, with a 120-second delay after each cycle to prevent overheating. After 3 min, if visual inspection revealed that the sample was insufficiently pulverised, an additional 60-second grinding with 30 sec in each cycle was performed. Subsequently, 1.2 mL of methanol (High-Performance Liquid Chromatography-grade; HPLC-grade) was added, and the tubes were incubated at RT for 24 h with slow shaking to extract the steroids (LLG uniROTATOR 2, LLG Labware, Germany). Following extraction, the samples were spun in a microcentrifuge (Centrifuge 5810 R; Eppendorf, Hamburg, Germany) at 12.000 rpm for 2 minutes, and 0.6 mL of each methanol extract was transferred to a 1.5 mL tube and evaporated using a vacuum evaporator (SPD1010 SpeedVac®, Thermo Fisher Scientific Inc., Waltham, MA, USA) at 45 °C and a vacuum of 0.6 torr. The dried extracts were reconstituted with 0.4 mL of the phosphate buffer provided in the assay kit (0 Calibrator DES6611CAL030 Lot 218 / 2026-04, Demeditec Diagnostics GmbH, Kiel, Germany). HCC was analysed in duplicate using a commercially available ELISA kit designed for free cortisol quantification in saliva (DES6611 Lot 218 / 2026-04, Demeditec Diagnostics GmbH, Kiel, Germany), following the manufacturer’s instructions. The resulting OD values were interpolated using GraphPad Prism software (version 4.01) and converted from ng mL⁻¹ to pg mg⁻¹ hair [45] for data analysis in R. The analytical sensitivity was 1.33 pg mg⁻¹ hair, based on the lowest standard provided in the assay kit. The intra- and interassay coefficients of variation were 5.0% and 7.3%, respectively.

### 2.4. Statistics

Statistical analyses and visualisation of Brix values (%), average daily weight gain (ADG) (kg), and HCC (pg mg⁻¹ hair) were conducted using RStudio (version 2026.07.1 + 147). The data evaluation started by defining an appropriate statistical model. The residuals were assumed to be normally distributed and homoscedastic. These assumptions were based on a graphical residual analysis.

Brix values were analysed using a linear model with cow–calf contact time (short vs. long) as the sole fixed effect. Differences between contact groups were evaluated using a one-way analysis of variance (ANOVA) based on F-tests.

ADG and HCC were analysed using a linear mixed-effects model (LMM) implemented with the *lmerTest* package. Cow–calf contact time (short vs. long), hutch type (single vs. pair), and their interaction were included as fixed effects. To account for the potential non-independence of calves housed together, housing unit identity was included as a random effect for pair-housed calves. Since individually housed calves did not share a housing unit, the random effect was restricted to pair housing, resulting in a partially nested model structure [46]. Both calves within each pair-housing unit belonged to the same cow–calf contact treatment group. The model was fitted using restricted maximum likelihood (REML) estimation. Fixed effects and their interaction were evaluated using Type III F-tests with Satterthwaite’s approximation for denominator degrees of freedom. Estimated marginal means (EMMs) and pairwise contrasts were calculated using the *emmeans* package. Four planned comparisons were performed: short versus long cow–calf contact within each hutch type, and single versus pair housing within each cow–calf contact time. Multiple contrast tests were performed following the principles described by Hothorn et al. [47] All comparisons were conducted using two-sided tests. To account for multiple testing, *p*-values and confidence intervals were adjusted simultaneously across the four contrasts using the multivariate t distribution (single-step adjustment), accounting for correlations among contrasts. A fixed random seed (12345) was used to ensure reproducibility of the numerical calculations. Statistically significant differences were declared at *p* < 0.05.

## 3. Results

In total, 35 experimental calves were sampled at day two of age and after weaning (88 ± 9 days of age). During the trial, 11 calves required medical treatment for minor clinical conditions, whereas 24 calves remained healthy throughout the study. Diagnoses included digestive problems (such as diarrhoea, coccidiosis and cryptosporidiosis), general signs of infection (fever), parasitic infestations, respiratory symptoms (coughing), and local inflammatory conditions (phlegmon and whitlow).

### 3.1. Immune status of the calves based on serum total protein

Adequate passive immune transfer via colostrum for animals with short and prolonged cow-calf-contact was verified by quantifying serum %Brix in the calves’ whole blood samples at 2 days of age by jugular venipuncture. 32 blood samples could be analysed in the laboratory. %Brix can be used as an indirect measure of passive immunity; it has a strong relationship with IgG because serum protein concentration is closely related to the amount of maternally derived IgG absorbed from colostrum [35]. There were no differences (*p* = 0.793) in %Brix between calves with short (S; n=15) and prolonged (L; n=17) contact-time with their dam, and all calves exhibited total serum protein concentrations indicative of successful passive immunity (S: 9.39 ± 1.29; L: 9.52 ± 1.35; mean ± SD). This prevented bias due to varying colostrum intake levels from affecting the results of the study and confirmed that there was no difference in health status between the treatment groups.

### 3.2. Average daily weight gain

The average daily weight gain (mean ADG, ± standard deviation; kg) for all calves was 0.850 kg ± 0.217 from housing in the calf hutch (1 or 2 days of age) to rehousing into groups (28 ± 5 days of age). The mean average daily weight gain from this period for each experimental group is shown in Table 3. The analysis of variance (ANOVA) revealed no effect of hutch type, cow-calf contact time or of an interaction (Table 4). Likewise, no differences between groups were found in pairwise comparisons with *p*-value adjustment (Supplementary Table S1).

**Table 3.** Average daily weight gain (ADG; kg) from 2 days of age to rehousing (28 ± 5 days of age) of the experimental groups.

| **Group** | **Mean ADG (kg)** | **± SD** | **n** | **min** | **max** |
|---|---|---|---|---|---|
| S1 | 0.948 | 0.209 | 12 | 0.553 | 1.285 |
| L1 | 0.856 | 0.148 | 13 | 0.548 | 1.064 |
| S2 | 0.738 | 0.322 | 5 | 0.445 | 1.104 |
| L2 | 0.715 | 0.212 | 5 | 0.217 | 0.917 |

S1 = short contact (< 12 h), single hutch; L1 = prolonged contact (24 – 48 h), single hutch; S2 = short contact (< 12 h), pair hutch; L2 = prolonged contact (24 – 48 h), pair hutch; Mean ADG = mean average daily weight gain in kg; SD = standard deviation; n = sample size; min = minimum, max = maximum.

**Table 4.** Two-way ANOVA table of average daily weight gain (ADG; kg) considering two factors with the interaction term (cow–calf contact × hutch type).

|  | **DF** | **Mean Square** | **f-Value** | ***p*-Value** |
|---|---|---|---|---|
| Cow–calf contact | 1 | 0.018 | 0.5714 | 0.460 |
| Hutch type | 1 | 0.073 | 2.2681 | 0.190 |
| Cow–calf contact : hutch type | 1 | 0.001 | 0.0434 | 0.837 |

DF = Degree of freedom

### 3.3. Hair cortisol concentration (HCC)

The concentration of hair cortisol was determined in pg mg⁻¹ of hair. HCC values ranged from 2.76 to 13.77 pg mg^-1^ of hair across all experimental groups (Table 5). The Supplementary Materials (Table S2) provide the HCC for each calf. The experimental period lasted 12 months, and thus, the calves were born and sampled in different seasons. Respective months of birth and sampling are shown in the Supplementary Materials (Table S2). The Analysis of Variance (ANOVA) revealed an effect of hutch type (*p* = 0.013), however, neither of cow-calf contact time nor of an interaction (Table 6). To further explore these differences, two-sided pairwise comparisons were conducted (Table 7). Four comparisons were performed: short versus long cow–calf contact within each hutch type, and single versus pair hutch within each cow-calf contact time. Pairwise comparisons revealed that S2-calves had lower HCC values than S1-calves (*p* = 0.047), while no difference (*p* = 0.891) was observed between L2- and L1-calves.

**Table 5.** Mean (± SD) hair cortisol concentrations (pg mg⁻¹ hair) by experimental group.

| Group | Mean | ± SD | min | max | n |
|---|---|---|---|---|---|
| S1 | 11.20 | 3.16 | 6.37 | 17.02 | 12 |
| L1 | 9.94 | 3.36 | 4.56 | 13.77 | 13 |
| S2 | 6.24 | 2.53 | 2.76 | 9.65 | 5 |
| L2 | 8.86 | 2.11 | 5.32 | 10.90 | 5 |

S1 = short contact (< 12 h), single hutch; L1 = prolonged contact (24 – 48 h), single hutch; S2 = short contact (< 12 h), pair hutch; L2 = prolonged contact (24 – 48 h), pair hutch; SD = standard deviation; n = sample size.

**Table 6.** Two-way ANOVA table considering two factors with the interaction term (cow–calf contact × hutch type).

|  | **DF** | **Mean Square** | **f-Value** | ***p*-Value** |
|---|---|---|---|---|
| Cow–calf contact | 1 | 3.338 | 0.358 | 0.554 |
| Hutch type | 1 | 65.170 | 6.985 | 0.013 |
| Cow–calf contact : Hutch type | 1 | 26.714 | 2.863 | 0.101 |

DF = Degree of freedom

**Table 7.** Pairwise comparisons (two-sided) of the mean hair cortisol concentrations in calves under different housing conditions and postpartum contact times.

| **Factor** | **Comparison** | **Estimate** | **SE** | **t-Value** | ***p*-Value** |
|---|---|---|---|---|---|
| 2 | L-S | 2.62 | 2.16 | 1.214 | 0.593 |
| 1 | L-S | -1.25 | 1.22 | -1.023 | 0.685 |
| L | 2-1 | -1.09 | 1.68 | -0.646 | 0.891 |
| S | 2-1 | -4.96 | 1.72 | -2.886 | 0.047 |

S = short contact (< 12 h); L = prolonged contact (24 – 48 h); 2 = pair hutch; 1 = single hutch; Estimate = estimate of the mean; SE = standard error; t-value = test statistic.

## 4. Discussion

Our working hypothesis was that early social contact would enhance calves’ capacity to cope with later management transitions, reflected by lower HCC values in S2- and L2-calves than in S1- and L1-calves. This hypothesis was only partially supported by the results. S2-calves exhibited lower HCC values during later management transitions, whereas L2-calves did not differ.

### 4.1. Immune status and colostrum supply

Newborn calves are highly dependent on the timely uptake of maternal immunoglobulins because immunoglobulins are not transferred from the dam to the foetus during gestation due to the epitheliochorial structure of the bovine placenta [48]. Consequently, calves are born with little to no circulating maternal antibodies and rely largely on colostrum intake shortly after birth to acquire passive immunity [48–51].

To benchmark the group-level status of passive immunity, our findings were compared against the herd-level standards proposed by Lombard et al. [52] with excellent (≥9.4% Brix), good (8.9–9.3% Brix), fair (8.1–8.8% Brix), or poor transfer of passive immunity (<8.1% Brix). Across experimental groups, mean serum Brix values ranged between 7.1% and 12.7%, aligning with typical field averages reported in large-scale studies [53, 54]. Brix values were comparable between all experimental groups, the newly introduced prolonged cow–calf contact period (24–48 h) or pair housing approach did not appear to negatively affect colostrum intake or the establishment of passive immunity in the calves.

### 4.2. Daily weight gain

Overall, the daily weight gains of calves were comparable across the experimental groups. This indicates that the newly introduced housing system, incorporating different forms of dam contact and socialisation in pair hutches, is generally feasible and supports comparable growth rates. This is contrary to previous studies, where social housing has been shown to increase feeding behaviour, feed intake, and activity compared with individual housing [59, 67]. Further research should investigate this relationship.

### 4.3. Hair cortisol concentration in calves

A significant main effect of hutch type on HCC was observed. However, the interaction between hutch type and cow-calf-contact was not significant. Consequently, S2-calves had lower HCC values than S1- calves.

Emotional bonding between calves and their dams begins shortly after birth and is influenced by the duration of contact [5, 55, 56]. Maternal affiliative behaviours, such as grooming and close proximity, may support the cow and calf [57, 58]. Early separation may therefore limit bond formation and affect subsequent stress responses. In our study, calves separated within 12 h of birth showed lower HCC values when subsequently pair-housed, whereas this pattern was absent after prolonged maternal contact. Pair housing may provide opportunities for affiliative interactions with a peer [59] and support coping with later challenges, such as rehousing and weaning [60].

Previous research suggests that social housing may reduce separation-related stress by providing social support [20, 61]. In our study, the presence of a second calf may have partially compensated for the loss of maternal contact but only when separation occurred immediately after birth. Although behaviour was not systematically assessed, qualitative observations of mutual grooming and close resting suggest affiliative interactions between paired calves. However, these observations provide only supportive, rather than direct, evidence of such social benefits.

Beyond stress-buffering effects, social housing may enhance calves’ cognitive abilities and behavioural flexibility and support more adaptive responses to novel situations [5, 62–66]. In our study, pair housing immediately after maternal separation may have partially compensated for the loss of maternal contact. Moreover, prior familiarity with at least one group member may have facilitated later social integration and reduced the stress associated with regrouping. Thus, social companionship may have provided benefits at both stages of rearing.

However, it remains unclear why calves that were separated from their dams after 24–48 h of contact (L1-, L2-calves) did not appear to benefit from this extended contact. It is also unclear whether prolonged dam contact may have attenuated or outweighed the positive effects of pair housing.

### 4.4. Limitations

#### 4.4.1. Comparability of hair cortisol concentrations with previous studies

A wide range of HCC values is reported in the literature (Table 8). As most studies sampled the back and shoulder region, this study’s results are consistent with previous findings in the literature, with values ranging from 2.4 to 52 pg mg⁻¹ across all calves and cattle. Differences between studies may be attributed to factors such as age, analytical methodology, and sampling site [36]. Initially, the measured values were consistently lower than those reported in most existing studies, prompting an investigation into the potential sources of this discrepancy. However, a recent study by Fushuku and Matsunaga [37] showed that values measured from hair in the chest region were significantly lower than those from hair on the back. Since the sampling site used in the present study was more comparable to the chest region, and many published values originate from methodological validation studies rather than comparisons between different husbandry systems, differences between studies should be interpreted with caution.

**Table 8.** Mean hair cortisol concentrations (pg mg⁻¹ hair) in calves and cattle reported in the literature.

| Sample Tool | Method | Concentration | Colour | Area | Age | Source |
|---|---|---|---|---|---|---|
| Scissors | EIA | ≈ 3.0 | NA | Tail tip | 30 d | [68] |
| Clippers | ELISA | 20.8 ± 3.0 | Black | Chest | ≈ 5.2 mo | [37] |
| NA | ELISA | 18.3 ± 2.8 | White | Chest | ≈ 5.2 mo | [37] |
| NA | ELISA | 0.036 | NA | Left shoulder | 16 wk | [67] |
| Clippers | (LC-MS/MS) | 2.4<br>(0.69–8.76) | Pigmented | Forehead | 156 ± 19 d | [69] |
| Electric hair clipper | ELISA | 19.6 ± 1.4 | Black | Caudodorsal | Cattle | [39] |
| Electric hair clipper | ELISA | 18.2 ± 1.4 | White | Caudodorsal | Cattle | [39] |
| Electric hair clipper | ELISA | ≈ 9.0 | Black | Back region | Cattle | [70] |
| Electric hair clipper | ELISA | ≈ 52.0 | Black and white | Shoulder, back, tail tip | Newborn | [29] |
| Electric hair clipper | ELISA | ≈ 6 | Black and white | Shoulder, back, tail tip | 6 mo | [29] |

\* Values reported in Knauer et al. [67] were recalculated from 0.97 ± 0.1 pg mg<sup>−1</sup> hair using the equation in Meyer et al. [45]. A = 0,97 pg/mL = 0,000097 µg/dL; B = 4.5 mg hair; C = 1 mL; D = 0,6 mL; E = 0,1 mL.

Total HCC in regrown hair collected in the current study may reflect chronic stress over about 2.7 months. Therefore, hair samples collected at 88 ± 9 days integrate cortisol during the period from rehousing into calf hutches to the early group housing phase, including weaning and regrouping as potential stressors. However, at this stage, it was not possible to distinguish which of these periods contributed most strongly to the observed cortisol levels. Consequently, the elevated or reduced hair cortisol levels observed at this stage cannot be attributed to a specific phase, as the measurements represent an integrated response over the entire period rather than phase-specific stress effects. Transferring calves to group housing is a significant stressor with presumed negative effects on health and growth [71].

#### 4.4.2. Lack of standardised HCC cut-off values limits comparability within and across housing systems

In this study, HCC was used as a small-scale indicator to assess system quality regarding animal welfare and stress [36]. However, the interpretation of HCC remains challenging due to the absence of validated cut-off values that define biologically relevant stress thresholds. However, interpreting HCC in calves remains challenging due to the lack of standardised reference values and variability caused by management, environment, individual history, and uncontrolled factors. Hair cortisol reflects cumulative hormonal exposure, but it may also include “background noise” from unknown or uncontrolled sources, which further obscures interpretation. Discontinuous observations resulted in so-called “black boxes”, which further limit the assessment of long-term stress. Given these limitations, there is a growing need for validated and robust welfare indicators to support reliable monitoring of system quality in scientific research and policy contexts [72].

#### 4.4.3. Implementation of pair housing as a new management practice on the study farm

Pair housing of calves was a new management approach on the study farm that had not previously been established in routine practice due to real-world implementation issues. We compared a long-established conventional calf management system (individual housing of calves; S1) to a newly introduced pair-housing approach (S2, L2). A major objective of the overall project was therefore to evaluate the implementation of this housing system under practical on-farm conditions and to assess its integration into a conventional dairy production system. Consequently, the study aimed to reflect real-life management conditions and their feedback on health-related outcomes. However, as the system was not yet fully established, systematic and random management-related effects may have contributed to the observed variability in the results.

Although pair housing may reduce stress in calves, its implementation can be challenging due to health issues, cross-suckling, and difficulties in forming age-matched pairs, particularly on small farms. Pairing calves of a similar age, preferably within ±1 day, may improve social compatibility and provide equal access to resources. Based on our observations in the present study, we tentatively suggest that pairing calves of a similar age, ideally within ±1 day, could benefit social compatibility and adaptation to pair housing. Larger farms may therefore be better suited to pair housing due to their larger herd sizes, which increases the probability of two calves being born within a one-day interval. However, this recommendation is preliminary, and further research should consider different management conditions and breed-related differences in growth and body size.

Separating cows and calves within 48 h is a simple and readily implementable management strategy that requires little additional investment, as existing farm infrastructure can be used. It may represent a practical compromise between immediate and prolonged cow-calf contact while addressing increasing animal welfare expectations. In Addition to these practical advantages, early separation may also offer economic benefits: early separation enables the efficient use of milk for human consumption and facilitates health monitoring [5, 13]. It is therefore hypothesised that this combination of practical and economic advantages may make the concept particularly attractive to farmers and encourage its adoption.

#### 4.4.3. Season bias considerations

Seasonal variation may have influenced the HCC values in our study. Fushuku and Matsunaga [37] reported higher HCC values in calves sampled in spring, suggesting that spring sampling may also have increased the HCC values in our study. Despite this, S2-calves sampled in spring showed significantly lower HCC values than the other groups. In contrast, L2-calves were predominantly sampled in spring and summer (Table S2), potentially introducing a seasonal bias that may have increased HCC values and masked lower values associated with pair housing. Balanced seasonal sampling is difficult under practical farm conditions due to farm size and seasonal calving patterns. Moreover, seasonal differences may be confounded by factors such as heat stress, respiratory infections, and changes in feed composition. Therefore, seasonal effects on HCC should be interpreted with caution.

## 5. Conclusions

Lower HCC values in S2 calves compared to S1 calves suggest potential differences in long-term cortisol exposure between hutch types. Early separation within 48 h may represent a practical and economically viable management strategy, balancing animal welfare considerations with farm efficiency. Furthermore, pairing calves of similar age, ideally within ±1 day, may improve social compatibility and facilitate adaptation to pair housing. However, HCC cannot be directly linked to specific management events or interpreted as a standalone welfare indicator. Larger, more balanced studies are therefore needed to clarify these interactions and evaluate HCC as an indicator of long-term physiological responses.

## Author Contributions

Conceptualisation, S.C.H.; formal analysis, I.K., S.C.H. and M.A.W.; investigation, I.K., M.A.W., and E.W.; resources, A.S. and S.C.H.; data curation, E.W. and I.K.; writing—original draft preparation, I.K.; writing— review and editing, A.S., M.S., and S.C.H.; supervision, M.S. and S.C.H.; project administration, A.S.; funding acquisition, S.C.H. All authors have read and agreed to the published version of this manuscript.

## Funding

This research was funded by the German Federal Ministry of Agriculture, Food and Regional Identity (BMLEH) and the Federal Office for Agriculture and Food (BLE): project “InnoRind - Zukunftsfähige Rinderhaltung in Deutschland unter Berücksichtigung von Tierwohl, Umweltwirkungen und gesellschaftlicher Akzeptanz”; Funding ID: 28N304205. I.K. received doctoral student funding from the H. Wilhelm Schaumann Foundation, Hamburg.

## Institutional Review Board Statement

The animal study protocol was approved by Kiel University animal welfare officer and the Schleswig-Holstein local authority (MLLEV-SH 64-12/23; March 18, 2025) for studies involving animals.

## Data Availability Statement

Data supporting this study are included within this article and/or supporting materials.

## Acknowledgments

The authors would like to thank our student assistants for their help with sample collection. We also thank a PhD colleague who coordinated the allocation of cows on the farm, as well as the farm staff for their support during the study. Additionally, we are grateful to the laboratory staff for performing the analyses. Finally, we want to thank Mario Hasler for his statistical advice. During the preparation of this manuscript, the authors used ChatGPT for language editing, including improvements to grammar, spelling, and phrasing. Further, ChatGPT was used to assist in coding during statistical analysis. The authors reviewed and edited the output and take full responsibility for the content of this publication.

## Conflicts of Interest

The authors declare no conflict of interest. The funders had no role in the design of the study; in the collection, analyses, or interpretation of data; in the writing of this manuscript; or in the decision to publish the results.

## Abbreviations

The following abbreviations are used in this manuscript:

HCC: hair cortisol concentration
ELISA: enzyme-linked immunosorbent assay
CCC: cow–calf contact
RT: room temperature
HPLC: High-Performance Liquid Chromatography
GLHT: linear hypothesis tests
ADG: average daily weight gains
ANOVA: Analysis of Variance

## Notes

### Competing Interest Statement

The authors have declared no competing interest.

