## Supplementary Table S1 for "The tell-tale hair: Effects of dam contact and hutch type on hair cortisol concentrations in female Holstein calves"

**Table S1.** Pairwise comparisons (two-sided) of the mean average daily weight gain (ADG; kg) in calves under different housing conditions and postpartum contact times.

| Factor | Comparison | Estimate | SE | t-ratio | *p*-Value |
|---|---|---|---|---|---|
| 2 | L-S | -0.0518 | 0.1990 | -0.261 | 0.9917 |
| 1 | L-S | -0.0913 | 0.0718 | -1.271 | 0.5328 |
| L | 2-1 | -0.1536 | 0.1640 | -0.935 | 0.7505 |
| S | 2-1 | -0.1931 | 0.1490 | -1.297 | 0.5384 |

S1= Short contact (< 12 h), single hutch; L1 = prolonged contact (24 – 48 h), single hutch; S2= short contact (< 12 h), pair hutch; L2 = prolonged contact (24 – 48 h), pair hutch; p-value adjustment: mvt method for 4 tests, Estimate = Estimate of the mean, SE = Standard Error, DF = Degree of freedom, t-ratio = test statistic, p-value.
